# CD29 enriches for cytotoxic human CD4^+^ T cells

**DOI:** 10.1101/2021.02.10.430576

**Authors:** Benoît P. Nicolet, Aurelie Guislain, Monika C. Wolkers

## Abstract

CD4^+^ T cell are key contributors in the induction of adaptive immune responses against pathogens. Even though CD4^+^ T cells are primarily classified as non-cytotoxic helper T cells, it has become appreciated that a subset of CD4^+^ T cells is cytotoxic. However, tools to identify these cytotoxic CD4^+^ T cells are lacking. We recently showed that CD29 (Integrin Beta 1, ITGB1) expression on human CD8^+^ T cells enriches for the most potent cytotoxic T cells. Here, we questioned whether CD29 expression also associates with cytotoxic CD4^+^ T cells. We show that human peripheral blood-derived CD29^hi^CD4^+^ T cells display a cytotoxic gene expression profile, which closely resembles that of CD29^hi^ cytotoxic CD8^+^ T cells. This CD29^hi^ cytotoxic phenotype was observed *ex vivo* and was maintained in *in vitro* cultures. CD29 expression enriched for CD4^+^ T cells, which effectively produced the pro-inflammatory cytokines IFN-γ, IL-2, and TNF-α, and cytotoxic molecules. Lastly, CD29-expressing CD4^+^ T cells transduced with a MART-1 specific TCR showed target cell killing *in vitro*. In conclusion, we here demonstrate that CD29 can be employed to enrich for cytotoxic human CD4^+^ T cells.

## INTRODUCTION

T cells are pivotal in mediating the clearance of pathogens, and in providing long-lasting immunity to recurring infections. Classically, T cells have been divided into cytotoxic CD8^+^ T cells and non-cytotoxic helper CD4^+^ T cells. Several studies have, however, demonstrated that the labor distribution of CD4^+^ and CD8^+^ T cells may be more nuanced. For instance, we and others demonstrated that human CD8^+^ T cells contain non-cytotoxic T cells (1–4). Furthermore, it has been documented that CD4^+^ T cells can be cytotoxic (5–9).

Cytotoxic CD4^+^ T cells can contribute to immune responses against viruses, such as Cytomegalovirus, Swine-flu, Epstein-Barr virus, and Human Immunodeficiency virus infections (10–14). They were also suggested to contribute to diseases such as IgG4-related disease (15). In addition, not only CD8^+^ T cells have been implicated in anti-tumor responses. Also cytotoxic CD4^+^ T cell responses were shown to substantially contribute to cancer clearance in mouse models (16, 17), and in human bladder cancer patients (6). Therefore, cytotoxic CD4^+^T cells could possibly be exploited for therapeutic purposes, such as cancer cellular immunotherapies. However, CD4^+^ T cells come in many flavors, and cytotoxic CD4^+^ T cells constitute only a subset of the CD4^+^ T cell pool. Furthermore, other CD4^+^ lineages such as regulatory T cells can hamper the anti-tumor potential (18). Therefore, preselection may be key for such approaches. Yet, the tools to do so are to date lacking.

We have recently shown that the expression of CD29 (Integrin β1; ITGB1) on human CD8^+^ T cells identifies cells that produce high amounts of cytotoxic molecules (i.e. Granulysin, Granzyme A and B, Perforin), of pro-inflammatory cytokines (i.e. IFN-γ and TNF-α) and of pro-inflammatory chemokines (i.e. CCL3, CCL4, CCL5) (1). Notably, the presence of CD29^hi^CD8^+^ T cells within tumors associated with better overall survival of melanoma patients, a feature that was also displayed in an increased *in vitro* killing capacity of tumor cells by CD29^hi^CD8^+^ T cells when compared to CD29^lo^ expressing CD8^+^ T cells (1). Whether the expression of CD29 is also linked to the cytotoxic function in human CD4^+^ T cells is not known.

In this study we show that human blood-derived CD29^hi^CD4^+^ T cells also display a cytotoxic gene expression profile, which closely resembles that of the CD29^hi^CD8^+^ T cell counterpart. CD29 expression enriched for CD4^+^ T cells that produced cytotoxic molecules, and the pro-inflammatory cytokines IFN-γ, IL-2, and TNF-α. This CD29^hi^cytotoxic phenotype of CD4^+^ T cells was observed *ex vivo*, and it was maintained upon T cell expansion *in vitro*. Lastly, we found that CD29-expressing CD4^+^ T cells were most potent in killing target cells. Therefore, we conclude that CD29 enriches for cytotoxic CD4^+^ T cells.

## MATERIALS AND METHOD

### Cell isolation and culture

Peripheral blood mononuclear cells (PBMCs) from de-identified healthy volunteers were isolated by Lymphoprep (density gradient separation; StemCell) and stored in liquid nitrogen until further use. The study was performed according to the Declaration of Helsinki (seventh revision, 2013). Written informed consent was obtained (Sanquin, Amsterdam, The Netherlands).

Cryo-preserved PBMCs were thawed and cultured in T cell medium (IMDM (LONZA) supplemented with 8% Fetal Calf Serum (FCS), 100U/mL Penicillin, 100µg/mL Streptomycin and 2mM L-glutamine). Cells were either directly used for flow-cytometry analysis, or were activated in T cell medium as previously described (19). Briefly, non-tissue culture treated 24-well plates (Corning, USA) were pre-coated overnight at 4°C with 4µg/mL rat α-mouse IgG2a (MW1483, Sanquin) in PBS. Plates were washed with PBS and coated for >2h with 1µg/mL αCD3 (clone Hit3a, eBioscience) at 37°C. 0.8×10^6^ PBMCs/well were seeded with 1µg/mL soluble αCD28 (clone CD28.2, eBioscience) in 1mL T cell medium. After 48h of incubation at 37°C, 5% CO_2_, cells were harvested, washed, and further cultured in standing T25/75 tissue culture flasks (Thermo-Scientific) at a density of 0.8×10^6^/mL, supplemented with 10ng/mL human recombinant IL-15 (Peprotech). Medium was refreshed every 5-7days.

### Flow-cytometry analysis

The cytokine production profile was determined by intracellular cytokine staining (ICCS) after T cell activation with 10ng/mL PMA and 1µM Ionomycin (Sigma-Aldrich), for 4h in the presence of 2µM monensin (eBioscience). Cells were stained in PBS + 1% FCS for live-dead marker (Invitrogen) and antibodies against: CD4 (RPA-T4, OKT4), CD8 (RPA-T8, SK1), CD29 (Mar4), CD38 (HIT2), CD27 (CLB-27), CD45RA (HI100) for 30min at 4°C in the dark. Cells were prepared with CytoFIX-CytoPerm kit following manufacturer’s protocol, and stained with antibodies against IFN-γ (4S.B3), IL-2 (MQ1-17H12), TNF-α (MAb11), Granulysin (DH2), Granzyme A (CB9), Granzyme B (GB11), Perforin (dG49). Cells were acquired on LSR II, Fortessa or Symphony (all BD) using FACS Diva v8.0.1 (BD). Data were analysed with FlowJo VX (TreeStar).

### Single cell RNA-sequencing analysis

Single cell RNA-sequencing (scRNA-seq) datasets were collected and re-analyzed from Guo et al. (20), Zheng et al (21), and Zhang et al (22). Count matrix was filtered for “PTH” and “PTY” to select peripheral blood CD4^+^ T cells expressing low and intermediate CD25, respectively. The Seurat R package (version 3.1.5; (23)) was used for scRNA-seq data analysis and batch-effect correction. Raw counts were log2-transformed, and used for further analysis. Differential expression (DE) analysis was performed, and significant genes were filtered for log2 fold change >0.25 and adjusted p-value <0.05. *ITGB1* (CD29) grouping was determined based on the “double peak” expression distribution of *ITGB1* in total CD4^+^ T cells (See Figure S1A, D). To determine T cell differentiation subsets, unbiased clustering was used. To attribute cell identities to clusters, differential gene expression analysis was performed on each cluster and compared to the rest of the CD4^+^ T cells (LFC>1 and p-adjusted < 0.05). Naïve-like clusters were characterized by high *CD27, CCR7, SELL, LEF1* and *TCF7* expression. Tcm-like cells were characterized by low amounts of cytotoxic molecules, and high expression of *CD27, CCR7*, and *SELL*. Tem and Teff were characterized by increasing expression level of cytotoxic molecules *GNLY, GZMB, GZMA, PRF1, SLAMF7* and *CX3CR1*. T cell clusters were then used to assess *ITGB1, SELL, CD27*, and *CX3CR1* expression.

### Generation of MART1-TCR expressing T cells

PBMCs from individual donors were activated for 48h with αCD3/αCD28 as described above, harvested and retrovirally transduced with the MART1-TCR, as previously described (1, 24). Briefly, non-tissue cultured treated 24 well plates were coated with 50µg/mL Retronectin (Takara) overnight and washed once with 1mL/well PBS. 300µL/well viral supernatant was added to the plate, and plates were centrifuged for 1h at 4°C at 2820 RCF. 0.5×10^6^ T cells were added per well, centrifuged for 10min at 180 RCF, and incubated overnight at 37°C. The following day, cells were harvested and cultured in T25 flasks at a concentration of 0.8×10^6^ cells/mL for 6-8 days in presence of 10ng/mL rhIL-15.

### Functional assays with MART1-TCR expressing T cells

The cytokine production of MART1-TCR transduced CD4^+^ T cells was determined by ICCS as described above. 100.000 TCR-transduced T cells (~80% TCR^+^) were co-cultured for 6h with 100.000 HLA-A2^+^ MART1^hi^ Mel 526 (MART1+), or HLA-A2^-^MART1^lo^ Mel 888 (MART1-) tumor cell lines. Cells were subsequently measured by flow cytometry, and populations of interest were gated and analyzed.

For killing assays, CD4^+^ MART1 TCR^+^ T cells were FACS-sorted for total CD4^+^, CD29^+^ (CD29^hi^) or CD29^-^CD38^+^ (CD29^lo^) expression, as previously described (1). Briefly, T cells were stained with antibodies against CD4, CD8, CD29, CD38, murine TCRβ (H57-597) and for live-dead marker in PBS + 1% FCS for 30min at 4°C. Cells were washed once with culture medium and sorted on a pre-cooled FACS Aria III (BD) sorter washed with 70% Ethanol. Sorted T cells were collected in culture medium, washed, and rested over-night in the incubator in medium at 37°C. Tumor cells were labelled with 1.5µM CFSE for 10 min at 37°C in FCS-free medium and washed 3 times with warm culture medium. 15×10^3^ tumor cells were co-cultured with MART1-TCR^+^FACS-sorted T cells for 20h, in a 3:1 ratio. Adherent and non-adherent cells were harvested from the wells to collect all remaining tumor cells. Dead tumor cells were quantified by flow-cytometry with Near IR live-dead marker on gated CFSE^+^ tumor cells.

### Statistical analysis and data visualization

Data generated with flow cytometry were analyzed with paired and ratio-paired t-test as indicated using GraphPad PRISM version 8. Differences were considered significant with a p-value < 0.05. Plots were generated with Seurat, ggplot2 version 3.0 and with GraphPad.

## RESULTS

### CD29 identifies human CD4^+^ T cells with cytotoxic gene expression

We first determined whether CD29 expression can identify cytotoxic CD4^+^ T cells, as we recently described for CD8^+^ T cells (1). To define the gene expression profile of CD29^hi^CD4^+^ T cells, we re-analyzed previously published single-cell RNA-sequencing (scRNA-seq) datasets of *ex vivo* blood-derived human CD4^+^ T cells (20–22). To exclude CD25^high^-expressing regulatory T cells from the analysis, we selected CD4^+^ T cells index-sorted for low and intermediate expression of CD25 (*see methods*), which yielded 3243 CD4^+^ T cells. When we analyzed the *ITGB1* (CD29) gene expression profile of these CD4^+^T cells, we observed a heterogenous distribution of *ITGB1* gene expression (Figure 1A). Nevertheless, we could define a clear cut-off of T cells with high (ITGB1^hi^) and low (ITGB1^lo^) *ITGB1* gene expression (Figure S1A). Differential gene expression analysis revealed that ITGB1^hi^CD4^+^ T cells preferentially expressed cytotoxic genes, including *GNLY* (Granulysin), *GZMA, GZMB, GZMH* (Granzyme A, B, and H), *PRF1* (Perforin), *FGFBP2, CCL5* (RANTES), and the cell surface markers *HLA-DR, NKG7* and *CX3CR1* (Figure 1B). This gene expression signature closely resembled that of previous studies on cytotoxic CD4^+^ T cells (7, 25, 26). In contrast, ITGB1^lo^CD4^+^ T cells showed higher gene expression of naïve and central memory-like markers such as *CCR7, CD27, SELL* (CD62L) and transcription factors *LEF1* and *SATB1*.

**Figure 1:**
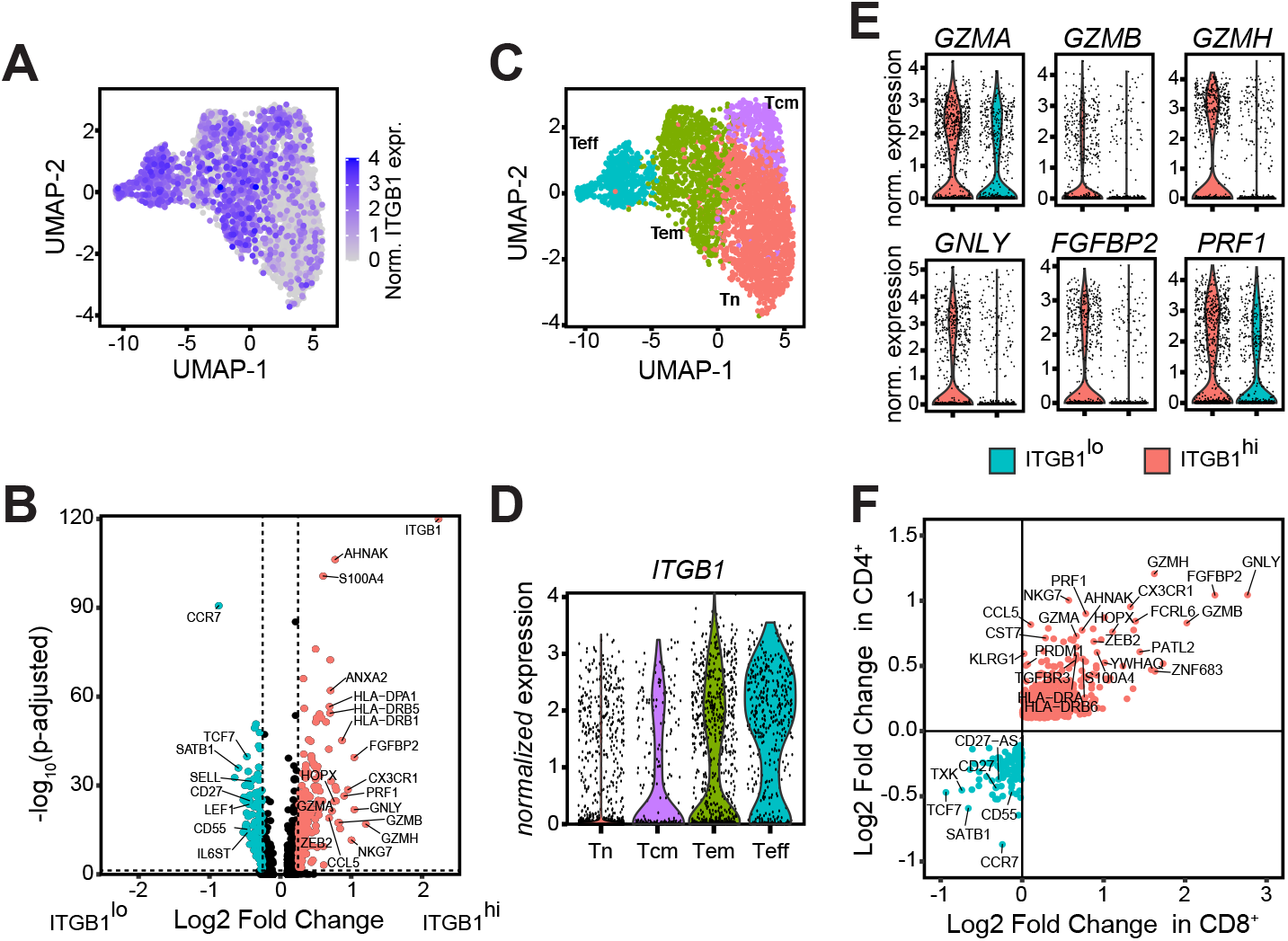
ITGB1 ^hi^ cells identifies human cytotoxic CD4^+^ T-cells in blood. Single-cell gene expression analysis re-analyzed from (20–22) of peripheral blood-derived FACS-sorted CD4^+^ T cells with low and intermediate CD25 protein expression. (**A**) UMAP projection of *ITGB1* (Integrin β1; CD29) gene expression in CD4^+^ T cells (n=3243, normalized expression). (**B**) Volcano plot of differentially expressed genes in CD4^+^ T cells with high (ITGB1^hi^) and low (ITGB1^lo^) ITGB1 expression (p-adj. <0.05 and log2 fold change > 0.25). Cut-off for *ITGB1* gene expression is depicted in Figure S1A. (**C**) Clusters and identity resulting from unsupervised clustering analysis of CD4^+^ T cell subsets from (A), Tn: Naïve; Tcm: central-memory; Tem: effector-memory; Teff: effector T cells. (**D**) Normalized *ITGB1* expression in CD4^+^ T cell subsets identified in (C). (**E**) Normalized gene expression for indicated genes in non-naïve CD4^+^ T cells, split according to *ITGB1* gene expression levels (see Figure S1D). (**F**) Comparison of differentially expressed genes and their corresponding log2 fold change between ITGB1^hi^ (red dots) and ITGB1^lo^ (blue dots) in CD4^+^ T cells from (B) and in CD8^+^ T cells from reference (1).

Peripheral-blood derived CD4^+^ T cells are comprised of T cells from distinct differentiation statuses, i.e. naïve (Tn), central-memory (Tcm), effector-memory (Tem) and effector T cells (Teff). We therefore asked how *ITGB1* gene expression was distributed over the different CD4^+^ T cell subsets. To this end, we used un-biased clustering and identified 4 differentiation clusters (Figure 1C, S1B; *see methods*). *ITGB1* gene expression was very limited on Tn cells (Figure 1D). Conversely, all non-naïve CD4^+^ T cell subsets contain T cells with high *ITGB1* gene expression, albeit to a different extent (Figure 1D). Measurements of CD29 protein expression on the different T cell subsets from blood-derived CD4^+^ T cells by flow-cytometry corroborated these findings (Figure S1C). To exclude that the cytotoxic gene expression profile of ITGB1^hi^CD4^+^ T cells was influenced by an over-representation of naïve (primarily ITGB1^lo^) T cells, we repeated the differential gene expression analysis on ITGB1^hi^ and ITGB1^lo^ non-naïve CD4^+^ T cells (Figure S1D, E). This analysis revealed a very similar gene expression profile of ITGB1^hi^ and ITGB1^lo^CD4^+^ T cells, with a high gene expression of *GZMA, GZMB, GZMH, GNLY, FGFBP2*, and *PRF1* on ITGB1^hi^CD4^+^ T cells, and on ITGB1^lo^CD4^+^ T cells a relatively high gene expression of *SATB1*, and of *CCR7* and *CD27* (Figure 1E, S1E, Table S1).

We next questioned how the cytotoxic gene expression profile of ITGB1^hi^CD4^+^ T cells related to that of ITGB1^hi^CD8^+^ T cells (1). We therefore compared the fold enrichment of differentially expressed genes of ITGB1^hi^ and ITGB1^lo^CD8^+^ T cells we previously described (1) with that of ITGB1^hi^ and ITGB1^lo^ CD4^+^ T cells. Remarkably, the cytotoxic gene expression features of ITGB1^hi^ CD8^+^ T cells was to a great extent shared with that of ITGB1^hi^CD4^+^ T cells (Figure 1F). This included the transcription factors Hobbit (*ZFN683;* (1, 25)), HOPX (27) and *ZEB2* (7, 28). We also found a set of genes that were commonly expressed by ITGB1^lo^ T cells, such as CCR7, SATB1, TCF7, CD55, the TCR-signaling molecule TXK, and the antisense long non-coding RNA *CD27-AS1*. Combined, our findings reveal that ITGB1^hi^CD4^+^ T cells have a cytotoxic gene expression signature that is shared with that of ITGB1^hi^CD8^+^ T cells.

### CD29 ^hi^ CD4^+^ T cells comprise cells that produce cytotoxic molecules and pro-inflammatory cytokines

We next questioned how the differential gene expression profile of ITGB1^hi^ and ITGB1^lo^CD4^+^ T cells translated into their protein expression profile. To test this, we first measured the protein expression of the cytotoxic molecules Granzyme A, Granzyme B, Granulysin, and Perforin in non-activated peripheral blood-derived CD4^+^ T cells by intracellular cytokine staining (ICCS). In line with the gene expression profile, CD29^hi^ (ITGB1^hi^) CD4^+^ T cells were the prime producers of all four cytotoxic molecules (Figure 2A, B).

**Figure 2:**
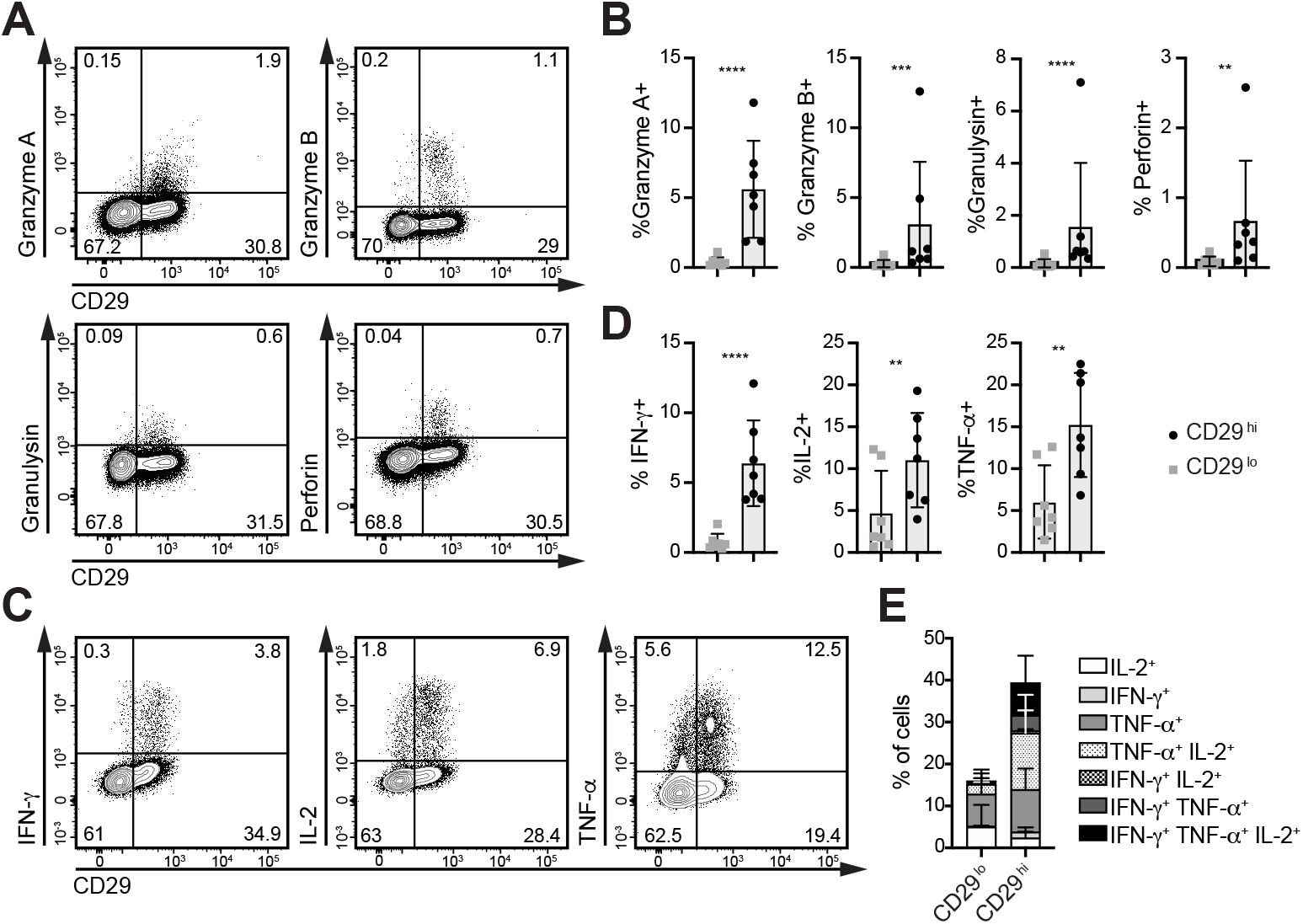
CD29 ^hi^ CD4^+^ T cells comprise cells that produce cytotoxic molecules and pro-inflammatory cytokines. (**A**) Representative dot plot of Granzyme A, Granzyme B, Granulysin and Perforin protein expression in resting peripheral blood-derived CD4^+^ T cells as determined by flow cytometry, and (**B**) compiled data of n= 7 donors. (**C**) Representative dot plot of IFN-γ, IL-2 and TNF-α protein production of peripheral blood-derived CD4^+^ T cells that were activated for 4h with PMA-Ionomycin, and (**D**) compiled data of n= 7 donors. (**E**) Fraction of IFN-γ, IL-2 or TNF-α (co-) producing CD29^lo^ and CD29^hi^CD4^+^ T cells. Differences between groups were determined with a ratio paired t-test. n.s.: not significant; ^**^: p<0.01; ^***^: p<0.001; ^****^: p<0.0001.

Not only the cytotoxic molecules, but also the pro-inflammatory key cytokines IFN-γ, TNF-α, and IL-2 are critical to the CD8^+^ T cells immune response (29–31). In CD8^+^ T cells, IFN-γ, TNF-α, and their co-expression of IL-2 strongly correlated with CD29 protein expression (1). To test whether CD29^hi^CD4^+^ T cells also display the preferential production of these three key cytokines, we stimulated blood-derived CD4^+^ T cells with PMA-ionomycin for 4h, and measured the production of IFN-γ, IL-2, and TNF-α protein by ICCS. As CD29^hi^CD8^+^ T cells alike, the percentage of CD29^hi^CD4^+^ T cells producing these three cytokines was substantially higher than that of CD29^lo^CD4^+^ T cells (Figure 2C, D). Notably, a substantial proportion of CD29^hi^CD4^+^ T cells comprised double and triple cytokine producers, a feature that is associated with the most potent T cell responses ((32, 33); Figure 2E)). Thus, CD29^hi^CD4^+^T cells include polyfunctional T cells that produce cytotoxic molecules and pro-inflammatory cytokines.

### In vitro cultured CD29^hi^ CD4^+^ T cells maintain their production profile

To determine whether the increased production of cytotoxic molecules and inflammatory cytokines by CD29^hi^CD4^+^ T cells was maintained upon T cell culture and expansion, we stimulated PBMCs with αCD3-αCD28 for 2 days, removed them from the stimulus, and expanded the cells for an additional 5 days in the presence of human recombinant IL-15. To separate CD29^hi^CD4^+^ T cells from CD29^lo^CD4^+^ T cells, we used the same gating strategy as previously employed for *in vitro* activated CD8^+^ T cells, i.e. CD29^hi^CD38^lo^(CD29^hi^) expressing T cells, and CD29^lo^CD38^hi^ (CD29^lo^) expressing T cells (Figure S2A, B; (1)). This gating strategy improved the distinction of the two T cell populations (Figure 3A, left panel).

**Figure 3:**
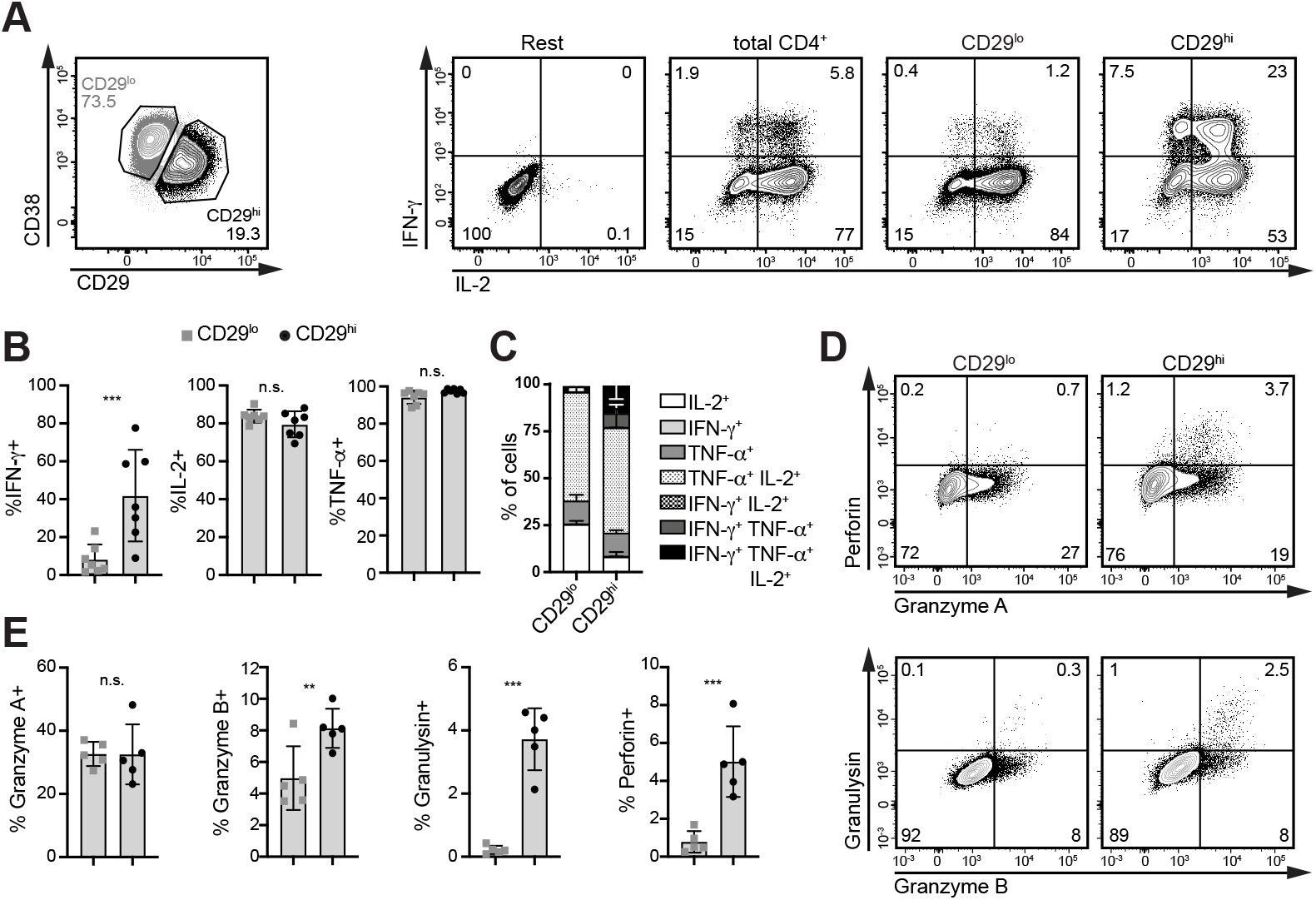
CD29^hi^ CD4^+^ T cells maintain their phenotype upon in vitro culture. PBMCs were activated for 2 days with αCD3-αCD28, removed from the stimulus and cultured for 4 days in recombinant human IL-15. (**A**) CD4^+^ T cells were assessed for CD29 and CD38 expression for the distinction of CD29^lo^ (CD29^low^ CD38^+^) and CD29^hi^(CD29^high^ CD38^-^; left panel) CD4^+^ T cells. Right panel: CD4^+^ T cells were re-stimulated with PMA-Ionomycin for 4h or left untreated (rest), and IFN-γ, IL-2 and TNF-α production was measured by ICCS. Representative dot plot, and (**B**) data compiled from 7 donors. (**C**) Distribution of single, double and triple cytokine-producing CD29^lo^and CD29^hi^CD4^+^ T cells. (**D**) Representative dot plot, and (**D**) quantification of Granzyme A, Granzyme B, Granulysin and Perforin expression in non-activated CD4^+^ T cells (n= 5 donors). Differences between groups were assessed using a ratio paired t-test. n.s.: not significant; ^*^: p<0.05; ^**^: p<0.01;^***^: p<0.001; ^****^: p<0.0001.

We next measured the cytokine production of the cultured CD4^+^ T cells by ICCS after T cell activation with PMA-ionomycin for 4h. Also upon T cell expansion *in vitro*, the percentage of IFN-γ producing T cells was significantly enriched in CD29^hi^CD4^+^ T cells compared to the total CD4^+^ T cell population, or to CD29^lo^CD4^+^ T cells (Figure 3A, B). Even though the percentage of IL-2 or TNF-α - producing cells was similar between CD29^lo^ and CD29^hi^CD4^+^ T cells, double or triple cytokine producers were enriched in CD29^hi^CD4^+^ T cells (Figure 3C), a feature that we also observed in CD29^hi^CD8^+^ T cells (1). In addition, while the percentage of TNF-α producing T cells was equally produced by CD29^lo^ and CD29^hi^CD4^+^ T cells (Figure 3B), the amount of cytokine produced per cell was higher in CD29^hi^CD4+ T cells as measured by the geometric Mean Fluorescence Intensity (geoMFI; Figure S2C).

When we analyzed the cytotoxic protein expression profile of non-activated *in vitro* cultured CD4^+^ T cells, we did not observe differences in the percentage of Granzyme A-expressing CD29^hi^CD4^+^ T cells compared to CD29^lo^CD4^+^ T cells (Figure 3 D, E). Yet, CD29^hi^CD4^+^ T cells showed higher Granzyme A protein expression per cell, as determined by the GeoMFI (Figure S2D). For Granzyme B, Perforin and Granulysin, however, we found substantial differences in non-activated CD4^+^ T cells (Figure 3D, E). In particular, Perforin and Granulysin were almost exclusively by non-activated CD29^hi^CD4^+^ T cells (Figure 3D, E). We thus conclude that CD29 expression enriches for CD4^+^ T cells with a cytotoxic phenotype also upon *in vitro* culture.

### CD29 ^hi^ CD4^+^T cells are superior IFN-γ and TNF-α producers in response to TCR activation

We next investigated the cytokine production profile of CD29^hi^CD4^+^ T cells in response to cognate antigen exposure. To this end, we retrovirally transduced CD4^+^ T cells with the codon-optimized MART1-TCR that recognizes the HLA-A^*^0201 restricted 26-35 epitope of MART1 (24, 34). Even though the MART1-TCR is MHC Class-I restricted, it was shown to elicit antigen-specific responses in MART1-TCR expressing CD4^+^ T cells (34).

The MART1-TCR transduction efficiency with ~80% was comparable between CD29^hi^ and CD29^lo^CD4^+^ T cells, and the TCR expression levels similar but slightly lower in CD29^hi^CD4^+^ T cells (Figure S2E). To determine the cytokine expression profile, MART1 TCR-engineered CD4^+^ T cells were co-cultured in a 1:1 ratio for 6h with HLA-A201^+^ MART1^high^-expressing melanoma tumor cell line (MART1+), or with HLA-A201^-^MART1^low^-expressing tumor cell line (MART1-(35, 36)). The cytokine production was measured by flow cytometry in CD29^hi^ and CD29^lo^CD4^+^ T cells (Figure 4A). The percentage of MART1 TCR-engineered CD29^hi^CD4^+^ T cells contained substantially more IFN-γ and TNF-α producing T cells in response to MART1+ tumor cells than the CD29^lo^CD4^+^ T cells (Figure 4A, B). The IL-2 production of CD29^hi^CD4^+^ T cells was similar to that of CD29^lo^CD4^+^ T cells. Notably, CD29^lo^CD4^+^ T cells comprised mostly IL-2 and TNF-α single producers, whereas most CD29^hi^CD4^+^ T cells produced IFN-γ, TNF-α and were enriched in IFN-γ^+^ TNF-α^+^ IL-2^+^ polyfunctional cells (Figure 4C). We detected Granulysin expression almost exclusively in resting MART1-TCR-expressing CD29^hi^CD4^+^ T cells (Figure 4D). Thus, MART1 TCR-expressing CD29^hi^CD4^+^ T cells maintain their cytokine production profile also when exposed to target cells.

**Figure 4:**
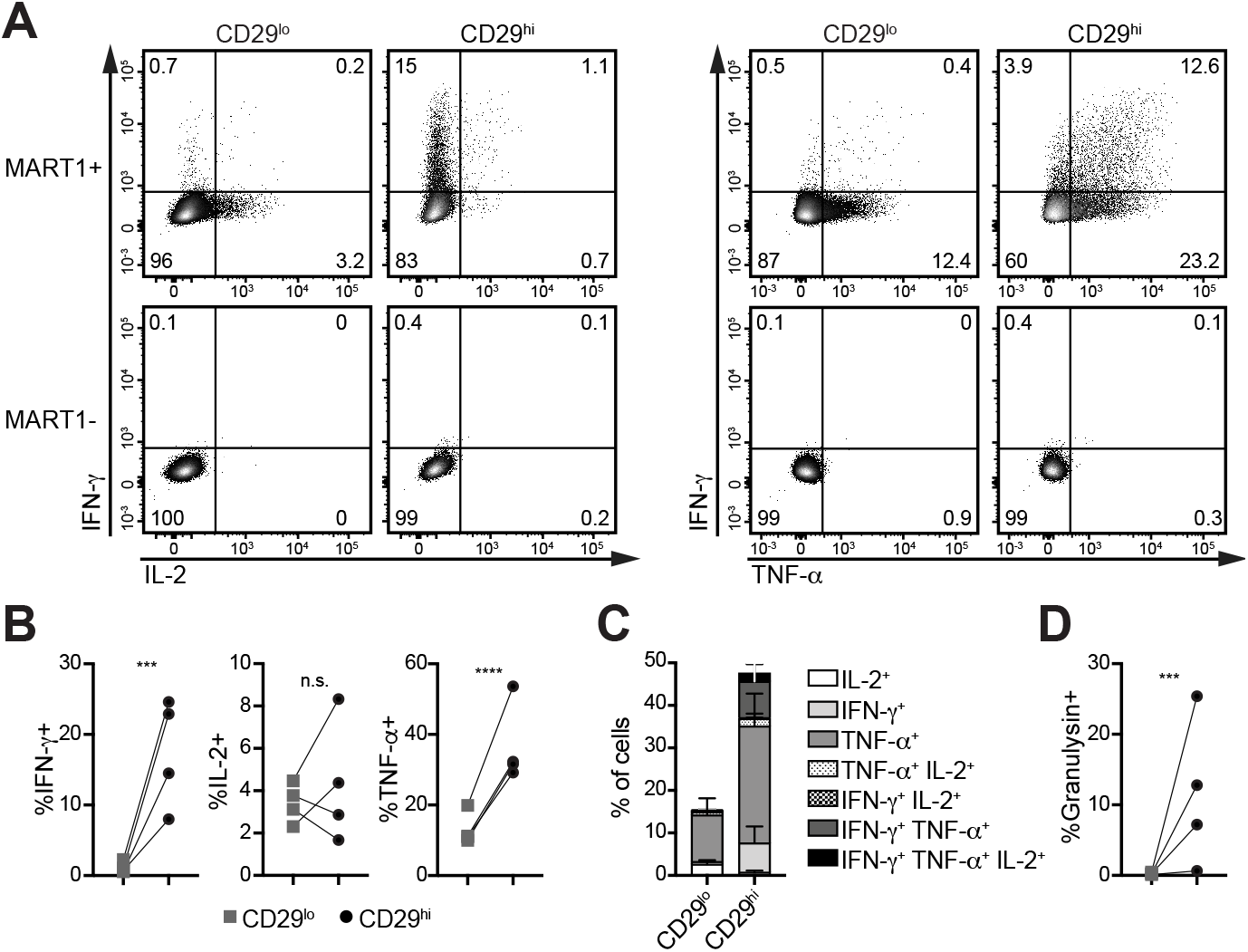
CD29^hi^ CD4^+^ T cells are superior IFN-γ and TNF-α producers *in vitro*. (**A**) MART1 TCR-engineered CD4^+^ T cells were cultured for 6h with MART1+ (upper panel) or MART1-(lower panel) tumor cells at a ratio of 1:1, and gated for CD29^lo^ or CD29^hi^ T cell populations. Representative dot plot of IFN-γ and IL-2 production (left panel) and of IFN-γ and TNF-α (right panel) as measured by ICCS. (**B**-**D**) Compiled data of 4 donors for (**B**) IFN-γ, IL-2 and TNF-α and (**C**) fraction of IFN-γ, IL-2, or TNF-α (co-) producing CD29^lo^ and CD29^hi^CD4^+^ T cells. (**D**) Granulysin content measured in resting CD29^lo^ or CD29^hi^ MART1 TCR-engineered CD4^+^ T cells. Differences between groups were assessed by ratio-paired t-test. n.s.: not significant; ^**^: p<0.01; ^***^: p<0.001; ^****^: p<0.0001.

### CD29^+^ enriches for cytotoxic CD4 T cells

Lastly, we determined the cytotoxic capacity of MART1 TCR-engineered CD29^hi^CD4^+^ T cells in response to tumor cells. CFSE-labelled MART1+ and MART1-tumor cells were co-cultured with FACS-sorted MART1 TCR-engineered total CD4^+^ T cells, or with CD29^lo^ or CD29^hi^ FACS-sorted TCR-engineered CD4^+^ T cells. After 20h, tumor cell killing was determined by live-dead marker labeling of all (adherent and non-adherent) tumor cells present in the co-culture.

The overall killing capacity of MART1 TCR-engineered total CD4^+^ T cells was low, but significant. MART1+ tumor cells, but not MART1-tumor cells, showed increased cell death in the presence of MART1 TCR-engineered T cells (Figure 5A, *left panel;* Figure 5B). Notably, in line with their cytotoxic gene and protein expression profile, FACS-sorted CD29^hi^CD4^+^ T cells from the same donors showed superior killing capacity when compared to CD29^lo^CD4^+^ T cells, or to total CD4^+^ T cells (Figure 5A, B). This augmented killing capacity of CD29^hi^CD4^+^ T cells was observed already at the low effector:target ratio of 3:1 that we used in our experiments (Figure 5A, B). Thus, selecting for CD29^hi^CD4^+^ T cells enriches for cytotoxic CD4^+^ T cells.

**Figure 5:**
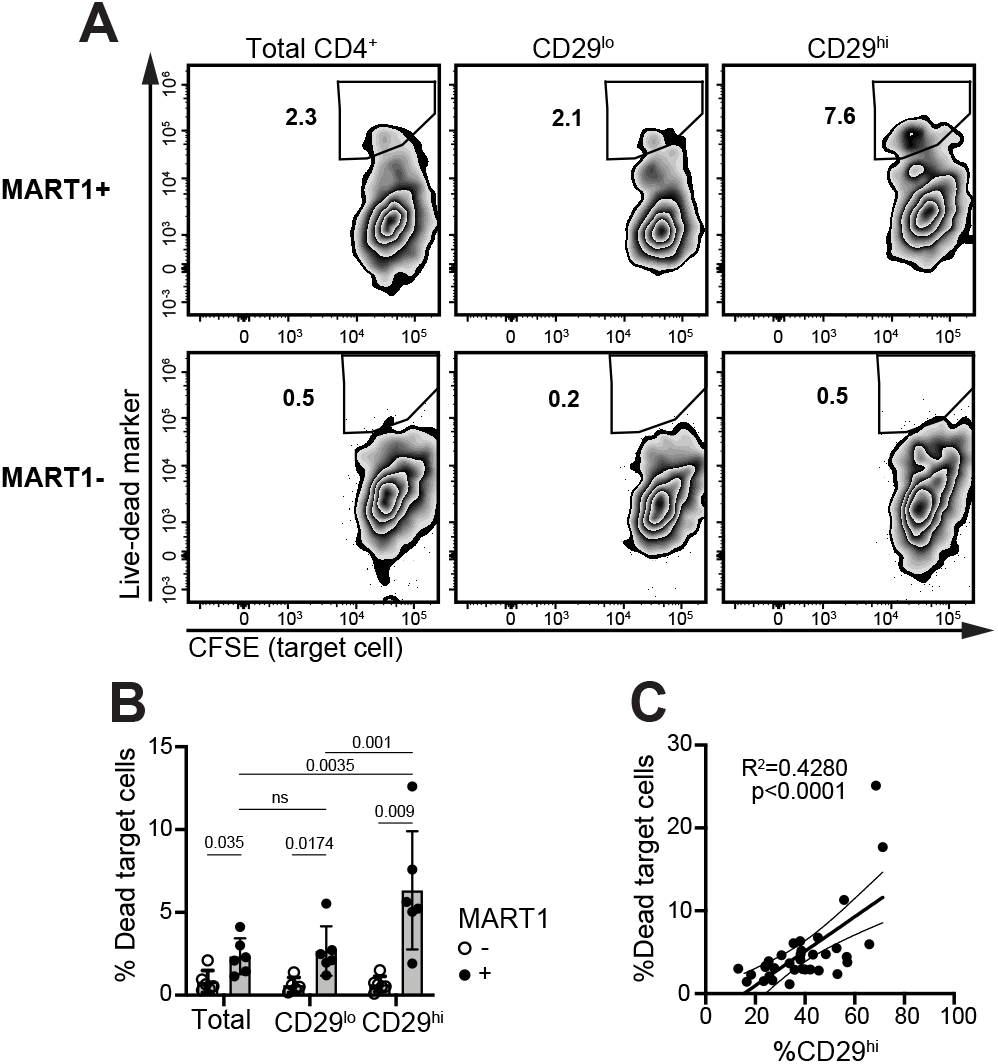
CD29 expression enriches for cytotoxic CD4^+^ T cells. (**A, B**) Representative flow-cytometry plots of MART1-TCR expressing CD4^+^ T cells (FACS-sorted total CD4^+^, and CD29^lo^ or CD29^hi^CD4^+^ T cells) were co-cultured for 20h with CFSE-labeled MART1+ (upper panel) or MART1-(lower panel) tumor cells at an Effector:Target ratio of 3:1. All cells (adherent and no-adherent cells were collected and dead tumor cells were determined by Near IR live-dead marker. (**A**) Representative dot plot, and (**B**) compiled data of 6 donors. Ratio-paired t-test per condition pairs, numbers in figure indicate p-values. (**C**) Correlation of tumor killing of total CD4^+^ MART1-TCR expressing T cells with the percentage of CD29^hi^ CD4^+^ T cells present in the T cell product (n=34, linear regression). n.s.: not significant.

We also defined how the overall CD29 expression levels of MART-1 TCR engineered CD4^+^ T cells related to their killing capacity. We therefore tested the tumor cell killing of MART-1 TCR engineered, FACS-sorted total CD4^+^ T cells from 34 individual donors. Importantly, we observed that the percentage of CD29-expressing CD4^+^ T cells strongly correlated with the killing capacity of the total CD4^+^ T cell population (p=<0.0001; Figure 5C). In conclusion, CD29-expressing CD4^+^ T cells comprise cytotoxic CD4^+^ T cells that display a polyfunctional cytokine production profile and that can effectively kill target cells.

## DISCUSSION

In this study, we show that CD29-expressing CD4^+^ T cells contain polyfunctional cytotoxic cells, a feature that is shared between CD4^+^ and CD8^+^ T cells. Our findings are concordant with another recent report on human cytotoxic T cells (26). Interestingly, the shared gene expression of the transcription factors ZNF683 (Hobit), HOPX and ZEB2 in CD29^hi^CD4^+^ and CD29^hi^CD8^+^ T cells suggests that cytotoxic CD4^+^ and CD8^+^ T cells share a transcriptional regulation network. In humans, Hobit is primarily found in effector CD8^+^ T cells, cytotoxic CD4^+^ T cells, and in resident memory T cells (Trm; (25, 37, 38)). Hobit is also known to repress CCR7, a gene we found associated with ITGB1^lo^CD4^+^ T cells (39). HOPX is involved in the regulation of granzyme B production in CD4^+^ T cells (27), and in the maintenance of effector memory within the type 1 helper T cells subset (40). Finally, the transcription ZEB2 works in coordination with T-bet to mediate the acquisition of cytotoxicity by CD8^+^ T cells (28). Differential gene expression of another transcription factor that was recently correlated with cytotoxic CD4^+^ T cells, thPOK, (27) was not found in ITGB1^hi^CD4^+^ T cells. Nevertheless, with our data combined, it is tempting to speculate that a transcription factor network involving ZFN683, ZEB2, HOPX, and possibly ThPOK are involved in the development or maintenance of CD29^hi^CD4^+^ and CD8^+^ T cells. Whether these transcription factors are involved in the expression of *ITGB1* is to date unresolved.

Also how CD29^hi^ cytotoxic CD4^+^ T cells are generated is to date not known. Cytotoxic CD4^+^ and their progenitor were recently described to be CD127^lo^ and CD127^hi^, respectively (7). Even though our scRNA-seq analysis did not reveal the CD127 (IL7R) gene to be differentially expressed in ITGB1^hi^ or ITGB1^lo^CD4^+^ T cells, CD29^hi^ T cells could very well arise from CD127^hi^ progenitors, and CD127 gene expression may redistribute upon differentiation.

Whether Integrin β1 has a role in the cytotoxic features or is a ‘bystander’ is yet to be determined. Integrin β1 can form heterodimers with 12 different alpha integrins (41). The composition the α-β integrin dimers confers ligand-specificity to the integrin complex. In T cells, the integrin complexes are important for cell-adhesion, cell migration, motility and signaling (42). For example, α4-β1 can act as a co-stimulatory molecule (43), as a receptor for CX3CL1 (fractalkine; (44)) and can skew CD4^+^ T cells towards a type 1 helper phenotype (45). It is also possible that CD29 expression contributes to T cell signaling and helps trigger cytotoxic functions, as observed in murine CD8^+^ T cell clones (46). Altogether, it is tempting to speculate that CD29 expression on cytotoxic CD4^+^ T cells is not merely a marker, but may also exert a function, for instance by providing additional signaling, or by stabilizing the cell-cell interaction.

Even though CD8^+^ T cells have been in the spotlight for therapies, recent findings suggest that cytotoxic CD4^+^ T cells are also involved in immune responses against tumors and viruses. In fact, the expansion of cytotoxic CD4^+^ T cells and their polyfunctionality is associated with healthy aging in super-centenarians (26), and they are shown to contribute to anti-viral responses (10–13). Furthermore, cytotoxic CD4^+^ T cells may substantially contribute to anti-tumor responses, as observed in bladder cancer patients (6). Whether these findings are applicable to other cancer types is not known yet and should be addressed. Nevertheless, in the context of cellular immunotherapies such as Chimeric Antigen Receptor T cells (CAR-T) and Tumor Infiltrating Lymphocytes (TILs) therapy, the selection of cytotoxic CD29-expressing CD4^+^ T cells could be of use to potentiate the efficacy of these therapies. Of note, the presence of cytotoxic CD4^+^ T cells may be beneficial in many, but not all cases. In fact, in SARS-Cov2 infected individuals, a higher number of cytotoxic CD4^+^ T cells was observed in severe compared to mild infections (47). This finding suggests that cytotoxic CD4^+^T cells contribute to the pathology in SARS-Cov2 infections, similar to what was previously observed in the IgG4-related disease (15).

In summary, we showed here that cytotoxic CD4^+^ T cells can be enriched for by the surface marker CD29, a feature that is shared with cytotoxic CD8^+^ T cells. Thereby, CD29 provides a robust marker to study human cytotoxic T cell responses.

## ACKNOWLEDGEMENTS

We would like to thank the blood donors for donation and E.Mul, M.Hoogenboezem and S.Tol for FACS sorting. We are grateful to B. Popovic, A. Jurgens, N. Zandhuis, S. Castenmiller for critical reading of this manuscript. We would like to thank R. Gomez-Eerland and T.N.M. Schumacher for providing the MART1 TCR system.

## AUTHOR CONTRIBUTIONS

B.P.N. and M.C.W. designed experiments and wrote the manuscript; B.P.N. and A.G. performed experiments and analyzed data; M.C.W. directed the study.

