## Supplemental figures for "CD29 enriches for cytotoxic human CD4^+^ T cells"

### Nicolet et al. Fig. S1

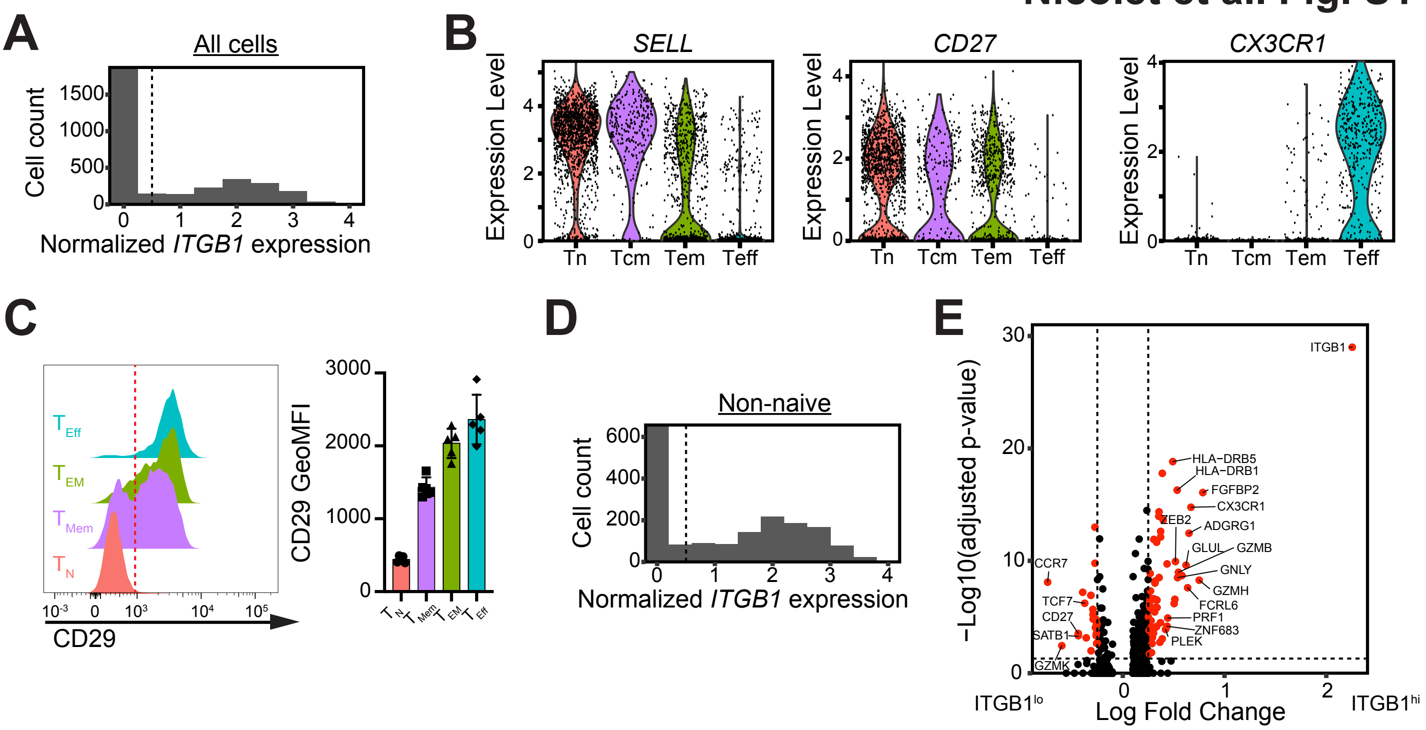


**Figure S1: ITGB1 gene expression on human blood-derived CD4$^+$ T-cells is associated with cytotoxic profile.** (**A**) Bar graph showing the expression of *ITGB1* (Integrin β1; CD29) of peripheral blood-derived CD4$^+$ T cells (n=3243, normalized expression) that were selected for low and intermediate level of CD25 gene expression (single cell RNA-sequencing data extracted from (18–20)). The dashed line depicts the expression cut-off used to define ITGB1$^{hi}$ and ITGB1$^{lo}$ cells based on *ITGB1* gene expression. (**B**) Normalized *SELL* (CD62L), *CD27*, and *CX3CR1* expression of different CD4$^+$ T cell differentiation subsets defined as in Figure 1C; naïve (Tn), central-memory (Tcm), effector-memory (Tem) and effector T cells (Teff). (**C**) Representative CD29 protein expression as determined by flow cytometry in Tn (CD45RA+ CD27+), Tmem (CD45RA- CD27+), Tem (CD45RA- CD27-), Teff (CD45RA+ CD27-) from blood-derived CD4$^+$ T cells (left panel). The dashed line represents the CD29 expression cutoff of CD29$^{lo}$ and CD29$^{hi}$ T cells. Compilation of CD29 geoMFI of all T cells in the 4 differentiation subsets (Tn, Tmem, Tem and Teff) of 5 donors (right panel). (**D**) Bar graph showing the expression of *ITGB1* gene expression in non-naïve CD4$^+$ T cells (n=1724; see methods; normalized expression). (**E**) Volcano plot depicting differentially expressed genes in non-naïve ITGB1$^{hi}$ and ITGB1$^{lo}$ CD4$^+$ T cells (log2 fold change >0.25 and adjusted p-value <0.05). Red color indicates genes considered differentially expressed.

Nicolet et al. Fig. S2

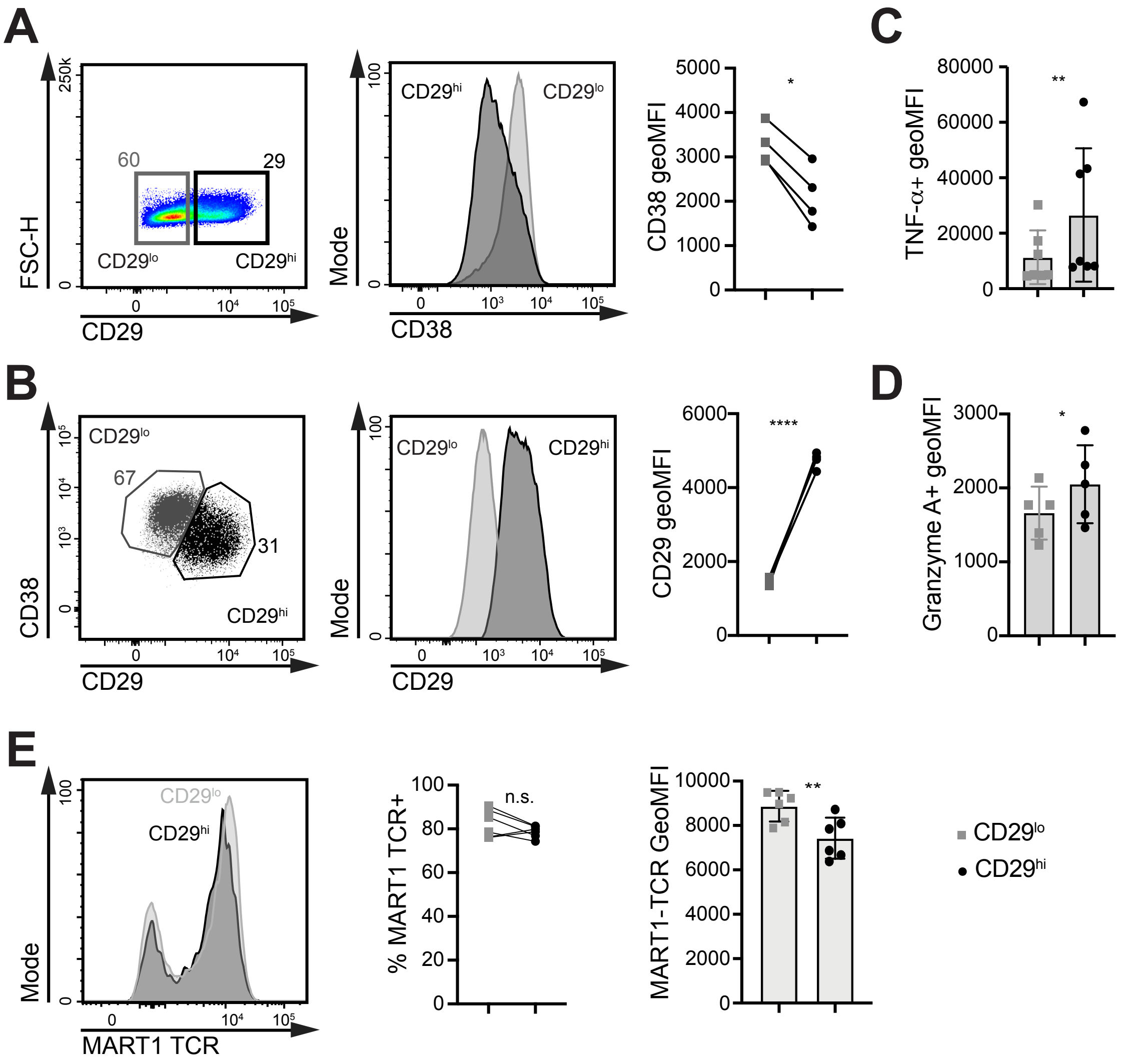


**Figure S2: CD29$^{hi}$ CD4$^{+}$ T cells maintain their phenotype upon *in vitro* expansion**

PBMCs were activated for 2 days with αCD3-αCD28 and rested for 4 days in hrIL-15. (**A**) Representative flow-cytometry plots of CD29 expression of CD4$^{+}$ T cells, and gating strategy used to define CD29$^{lo}$ and CD29$^{hi}$ CD4$^{+}$ T cells (left panel). Representative histogram of CD38 expression of CD29$^{lo}$ and CD29$^{hi}$ CD4$^{+}$ T cells (middle panel) and compiled geometric mean fluorescence intensity (geoMFI) expression of 4 donors (right panel). (**B**) Gating strategy (left panel), representative CD29 expression (middle panel), and compiled geoMFI (right panel) of CD29$^{lo}$ (CD29$^{low}$ CD38$^{+}$) and CD29$^{hi}$ (CD29$^{high}$ CD38$^{-}$) CD4$^{+}$ T cells. (**C**) GeoMFI of TNF-α$^{+}$ producers of CD29$^{lo}$ and CD29$^{hi}$ CD4$^{+}$ T cells activated with PMA-Ionomycin for 4h, from Figure 3C (n=7 donors). (**D**) GeoMFI of granzyme A$^{+}$ producers from Figure 3E, measured in non-activated CD4$^{+}$ T cells (n=5 donors). (**E**) CD4$^{+}$ T cells were retrovirally transduced with the codon-optimized MART1-TCR. MART1-TCR transduction efficiency was assessed by flow-cytometry at day 7 post transduction. Representative histogram of MART1-TCR expression (left panel), the percentage of MART1-TCR$^{+}$ (center panel), and the GeoMFI of MART1-TCR$^{+}$ in CD29$^{lo}$ and CD29$^{hi}$ CD4$^{+}$ T cells (right panel) of 6 donors. Differences between groups were determined with a ratio paired t-test. n.s.: not significant; *: $p<0.05$; **: $p<0.01$; ****: $p<0.0001$.
